# Quantitative analysis of neuronal mitochondrial movement reveals patterns resulting from neurotoxicity of rotenone and 6-hydroxydopamine

**DOI:** 10.1101/2021.02.24.432715

**Authors:** Rui F. Simões, Rute Pino, Maurício Moreira-Soares, Jaromira Kovarova, Jiri Neuzil, Rui Travasso, Paulo J. Oliveira, Teresa Cunha-Oliveira, Francisco B. Pereira

## Abstract

Alterations in mitochondrial dynamics, including their trafficking, can present early manifestation of neuronal degeneration. However, current methodologies used to study mitochondrial trafficking events rely on parameters that are mostly altered in later stages of neurodegeneration. Our objective was to establish a reliable computational methodology to detect early alterations in neuronal mitochondrial trafficking. We propose a novel quantitative analysis of mitochondria trajectories based on innovative movement descriptors, including straightness, efficiency, anisotropy, and kurtosis. Using biological data from differentiated SH-SY5Y cells treated with mitochondrial toxicants 6-hydroxydopamine and rotenone, we evaluated time and dose-dependent alterations in trajectory descriptors. Mitochondrial movement was analyzed by total internal reflection fluorescence microscopy followed by computer modelling to describe the process. The stacks of individual images were analyzed by an open source MATLAB algorithm (www.github.com/kandelj/MitoSPT) and to characterize mitochondria trajectories, we used the Python package trajpy (https://github.com/ocbe-uio/trajpy/). Our results confirm that this computational approach is effective and accurate in order to study mitochondrial motility and trajectories in the context of healthy and diseased neurons in different stages.

## 1. Introduction

Neurons are polarized post-mitotic cells encompassing three structurally, functionally, and metabolically distinct domains, i.e. the cell body, dendrites with numerous branches, and the axon. These domains display unique metabolic and energetic needs, and they rely on mitochondrial adenosine triphosphate (ATP) production to accomplish their specific functions (1–3). Mitochondria-produced ATP is vital for neuronal cell growth and survival, synapse formation and assembly, generation of action potentials, synaptic transmission and synaptic vesicle trafficking (4–6). Additionally, mitochondria are also pivotal in calcium (Ca^2+^) homeostasis in neuronal cells, buffering transient Ca^2+^ levels by its sequestration and release, as needed (7, 8). As individual neuronal domains feature specific needs for the level of Ca^2+^ as well as metabolites, their homeostasis is maintained by corresponding number of mitochondria (1, 9, 10).

Due to their morphological and metabolic characteristics, neuronal cells have developed mechanisms to transport mitochondria along microtubular tracks. The movement from the cell body to cellular extremities (anterograde transport) is mediated by the kinesin-1 family proteins, while dynein proteins are responsible for the opposite movement (retrograde transport). Both types of transport are dependent on ATP hydrolysis (11, 12). Movement of mitochondria is dependent on the polarity of microtubules, polymeric structures composed of α- and β- tubulin, that polymerize from the minus to the plus end. In axons, the minus end is directed towards the cell body and the plus end to the cell extremity (13, 14). Thus, kinesins carry mitochondria from the minus to the plus end and dyneins from the plus to the minus end (15).

Mitochondrial trafficking is also dependent on adaptor proteins, which ensure targeted and efficient transport regulation. The trafficking kinesin-binding (TRAK) proteins 1 and 2 bridge the mitochondrial rho (MIRO) 1 and 2 proteins and kinesins to control mitochondrial anterograde trafficking (16, 17). Relevant for the retrograde transport, dynactin binds to dynein and to the microtubules, enhancing dynein motor processivity (11, 18). Mitochondrial docking processes allow mitochondria to remain stationary in areas with elevated ATP demand and Ca^2+^ buffering dependency (1). It has been described that between 10% and 40% of mitochondria in a neuronal cell are in motion, while 60% to 90% of the organelles are stationary (10, 19, 20).

Since mitochondria are physically allocated to areas with higher metabolic activity and also based on regulation of Ca^2+^ homeostasis, aberrations in mitochondrial dynamics, metabolism and mobility, leading to altered ATP production and lower Ca^2+^ buffering capacity, are involved in the development of neurodegenerative pathologies, such as Alzheimer’s disease, Huntington’s disease, Parkinson’s disease, and amyotrophic lateral sclerosis (1). Furthermore, it was previously shown that alterations in mitochondrial motility appear prior to the first signs of neurodegeneration (such as degeneration of axons and cell death) (21, 22).

Mitochondria move along short or long paths with varying velocities and directions, often altering those parameters as a response to different stimuli (23). Additionally, mitochondria undergo morphological alterations during movement, posing an increased challenge in the identification and segmentation of individual mitochondria. A further obstacle is the low signal-to-noise ratio of many microscopic approaches, yielding poor quality images (24). Associated video acquisition processes can be detrimental due to photobleaching and phototoxicity (25). Therefore, the videos taken under these microscopy approaches are either short (2-5 min) with 1-2 frames per second (fps) (20, 24, 26) or longer (30 min) but with 1 frame every 5-10 s (27). Short videos cannot collect all features required to characterize mitochondrial motion while in longer movies, important information may be lost between frames.

To circumvent the limitations described, we used here total internal reflection fluorescence (TIRF) microscopy that takes advantage of a special mode of sample illumination, exciting only fluorophores located near the sample interface (about 100 nm), without exciting sample regions located further away. Images obtained with this microscopy technique present higher signal-to-noise ratio and almost nonexistent out-of-focus fluorescence, preventing photobleaching and phototoxicity (28–30).

We exposed the cells to mitochondrial toxins, evaluated the mitochondrial movement by TIRF microscopy, and used computer modelling to describe the process. Our results present a new quantitative paradigm of mitochondrial dynamics in health and diseased neuronal cells.

## 2. Results

### 2.1. 6-OHDA and rotenone decreased ATP levels in a concentration and time-dependent manner

Mitochondrial trafficking in neuronal cells is highly dependent on ATP consumption, since kinesin and dynein transport requires ATP hydrolysis (Hirokawa et al. 2010). We initially measured cellular ATP levels after treating cells for 24 h and 96 h with 6-OHDA (Fig 1 a) and rotenone (Fig 1). While there was little if any effect of the agents at 24 h, 96h-treatment caused considerable decrease in ATP levels (Fig 1).

**Fig 1.**
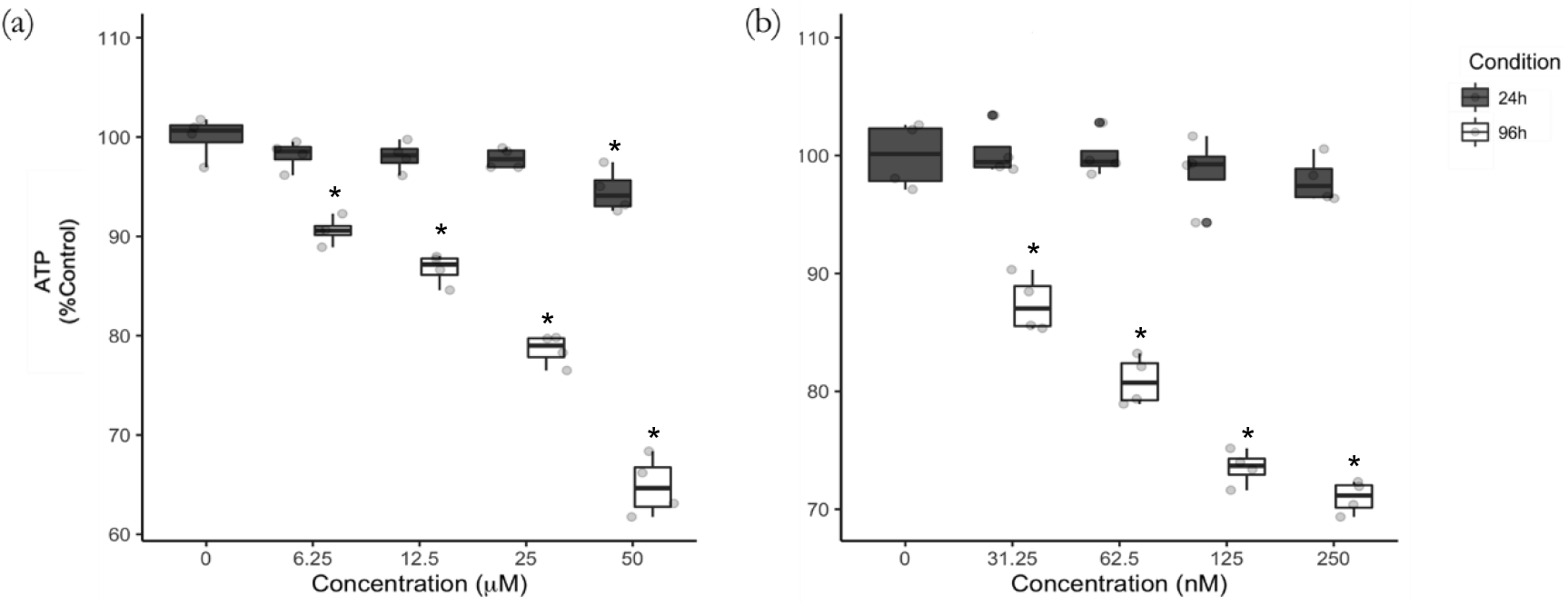
Treatment with 6-OHDA (a) and rotenone (b) for 24h and 96 h causes different decreases of ATP levels. Differentiated SH-SY5Y cells were treated with 6-OHDA (a) or rotenone (b) for 24 and 96 h. Data are presented as boxplots, in which each dot represents an independent cell population (n=4), in duplicate. Kruskal-Wallis test (one-way ANOVA on ranks) pairwise (control vs 6-OHDA or control vs rotenone) was used to assess statistical significance, (*) p< 0.05.

### 2.2. 6-OHDA and rotenone reduced the level of βIII tubulin

We next tested the effect of 6-OHDA and rotenone on tubulin levels in differentiated SH-SY5Y cells. All 6-OHDA treatments (except 12.5 μM for 96 h) significantly decreased the levels of tubulin (Fig 2 a and b). Regarding rotenone, 125 nM and 250 nM for the 24-h time point induced an evident decreased in tubulin levels (Fig 2 c). This effect was considerably stronger for the 96-h treatment (Fig 2 d). Representative images (Fig 2 e and f) of cells treated with different concentrations of 6-OHDA (ii-v) and rotenone (vi-ix) for 24 h (e) and 96 h (f) are presented below.

**Fig 2.**
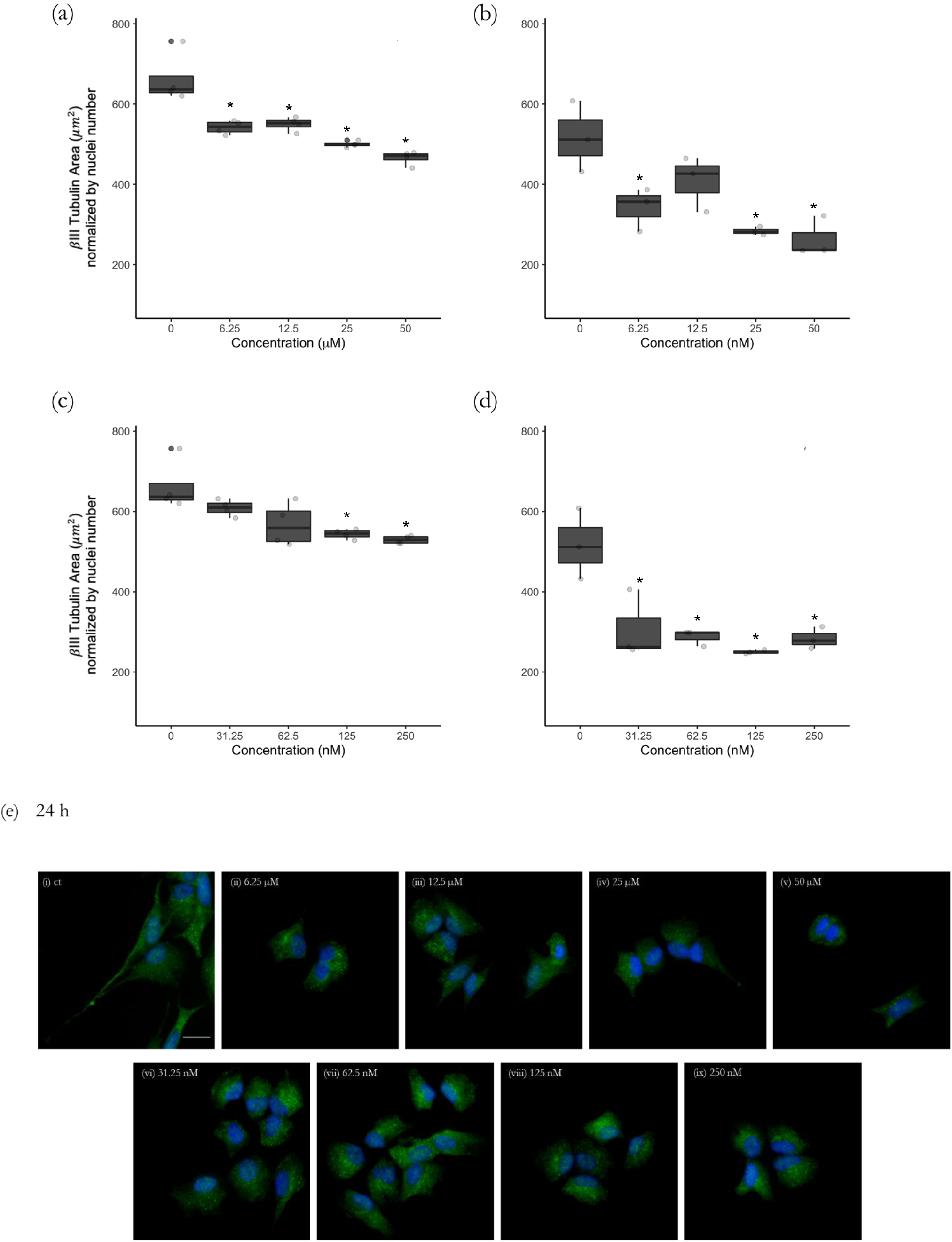
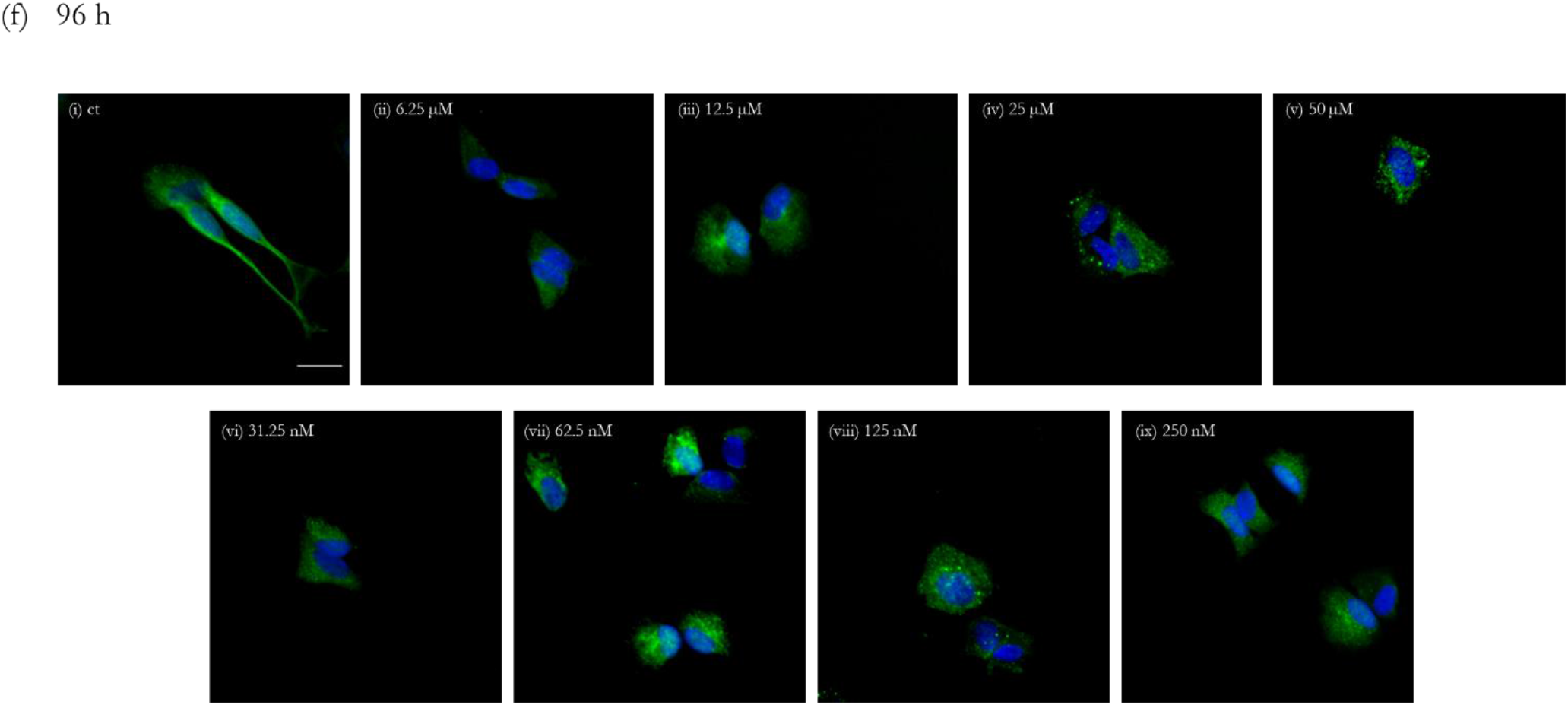
Incubations with 6-OHDA and rotenone decreases the level of βIII tubulin in a dose-dependent manner. Incubations with 6-OHDA for 24 h (a) and 96 h (b) or rotenone for 24 h (c) and 96 h (d) induced a significant decrease in βIII tubulin level. Data are presented as boxplots, in which each dot represents an independent cell population (n=4) in duplicate. Kruskal-Wallis test (one-way ANOVA on ranks) pair-wise (control vs 6-OHDA or control vs rotenone) was used to assess statistical significance, (*) p< 0.05. Immunofluorescence images of βIII tubulin in differentiated SH-SY5Y cells were acquired using a 20x objective and the IN Cell Analyzer 2200. Scale bar = 20 μm. Nuclei staining is presented in blue and βIII tubulin is presented in green (e and f). Cells were treated for 24 h (e) or 96 h (f) with 6.25 (ii), 12.5 (iii), 25 (iv) and 50 μM (v) of 6-OHDA or with 31.25 (vi), 62.5 (vii), 125 (viii) and 250 nM (ix) rotenone. Non-treated cells are presented in part (i) of both (e) and (f) panels.

### 2.3. Mitochondrial net displacement is decreased by treatment with 6-OHDA and rotenone

Treatment with 50 μM 6-OHDA for 24 h resulted in significantly smaller mitochondrial net displacement. When incubated for 96 h, all 6-OHDA concentrations substantially decreased mitochondrial net displacement in differentiated SH-SY5Y cells when compared to their control counterparts (Fig 3 a). Incubation with 62.5 nM rotenone for 24 h resulted, on average, in a 22% reduction of mitochondria net displacement, reaching statistical significance at 125 and 250 nM. Cells treated for 96 h with rotenone presented a significant decrease in mitochondrial net displacement when compared to untreated counterparts (Fig 3 b).

**Fig 3.**
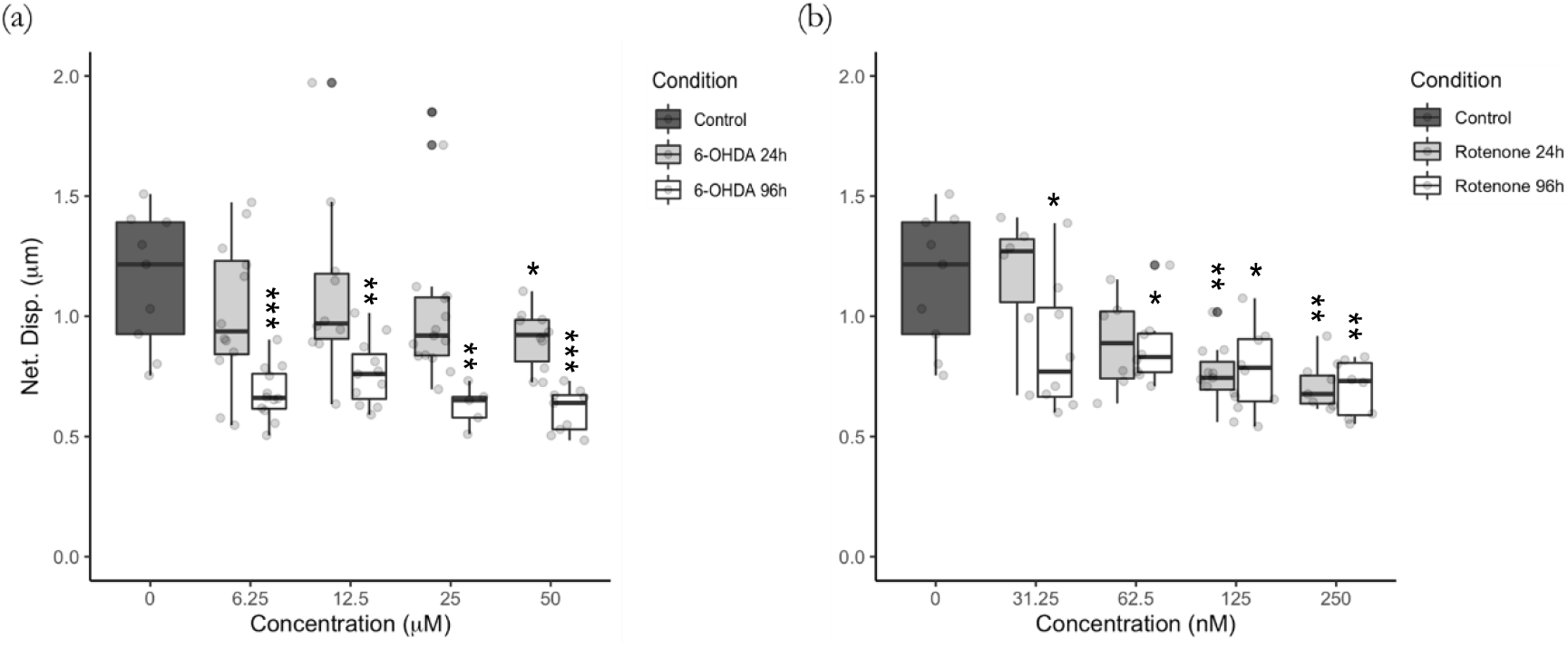
6-OHDA (a) and rotenone (b) reduced mitochondrial net displacement. Mitochondria were labeled with the fluorescent dye MitoTracker Red CMXRos, their movement followed, and trajectory net displacement was calculated as stated in Materials and Methods. Data are presented as boxplots, in which each dot represents the mean of each mitochondrial movement per video frame (n=5 to 15). Kruskal-Wallis test (one-way ANOVA on ranks) pair-wise (control vs 6-OHDA or control vs rotenone) was used to assess statistical significance, (***) p< 0.001 (**), p< 0.01, (*) p< 0.05.

### 2.4. Mitochondrial mean velocity is decreased by treatment with 6-OHDA and rotenone

Our results indicated that mitochondria move in control cells with the rate of 0.1 to 0.6 μm/s. (Fig 4 a). Differentiated SH-SY5Y cells treated with 6-OHDA for 24 h exhibited a significant decrease in mitochondria mean velocity when incubated with 25 μM and 50 μM. Cells incubated with 12.5 μM, 25 μM and 50 μM 6-OHDA for 96 h showed mitochondrial movement, on average, 55% slower than mitochondria in untreated cells, reaching statistical significance when treated with 6.25 μM (Fig 4 a). Rotenone-treated cells incubated with 125 nM and 250 nM for 96 h revealed 55% slowed movement of mitochondria, although this did not reach significance (Fig 4 b).

**Fig 4.**
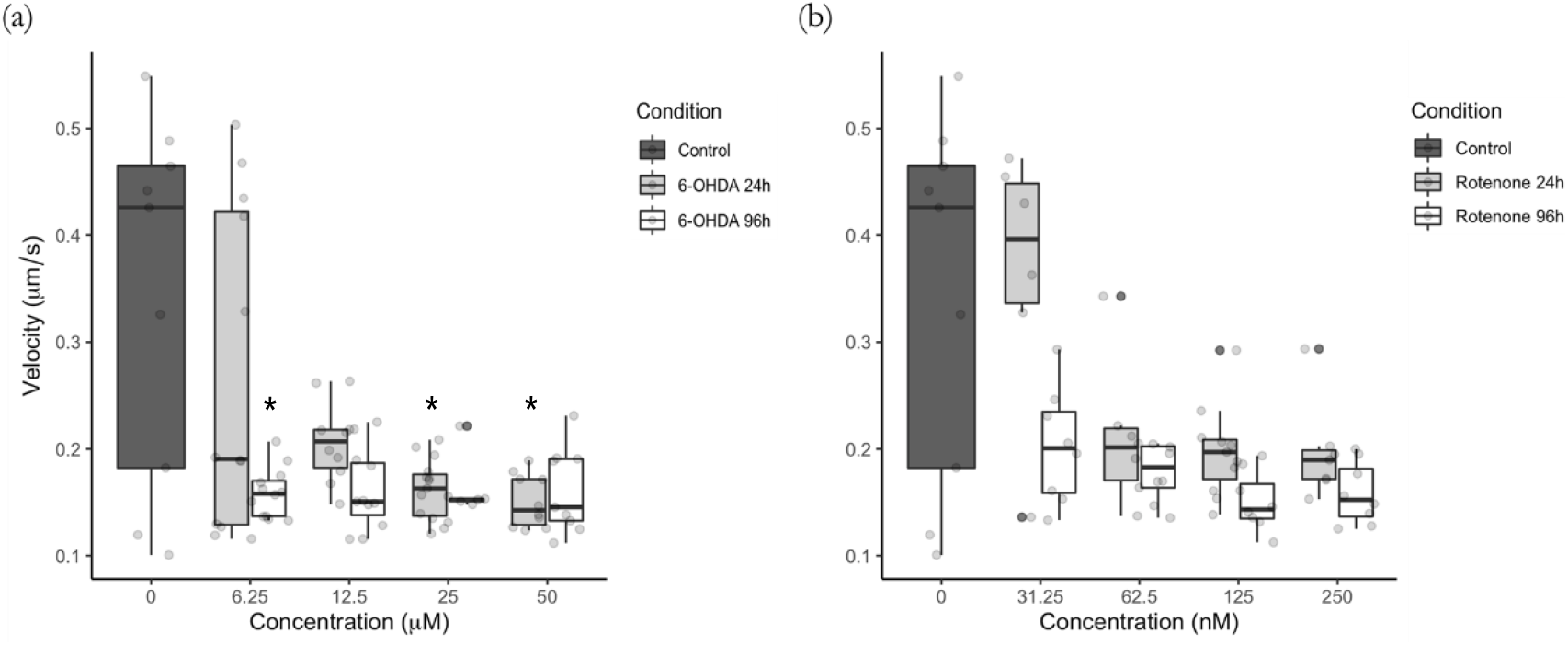
Mitochondrial mean velocity is lower due to 6-OHDA (a) and rotenone (b) treatment. Mitochondria were labeled with the fluorescent dye MitoTracker Red CMXRos, their movement followed, and trajectory mean velocity was calculated as stated in Materials and Methods. Data are presented as boxplots in which each dot represents the mean of each mitochondrial movement per video frame (n=5 to 15). Kruskal-Wallis test (One-way ANOVA on ranks) pair-wise (control vs 6-OHDA or control vs rotenone) was used to assess statistical significance, (*) p< 0.05.

### 2.5. Mitochondrial movement trajectory is affected by 6-OHDA and rotenone

Concerning mitochondria trajectory straightness, mitochondria in cells treated for 24 h with 50 μM 6-OHDA showed non-straight movement trajectories when compared to control cells. The other concentrations of 6-OHDA caused only minor alteration of mitochondrial movement trajectories (Fig 5 a). Rotenone at 125 nM induced a 17% decrease in mitochondria trajectory straightness although this was not significantly different from parental cells. No alterations were found for the other rotenone concentrations (Fig 5 b).

**Fig 5.**
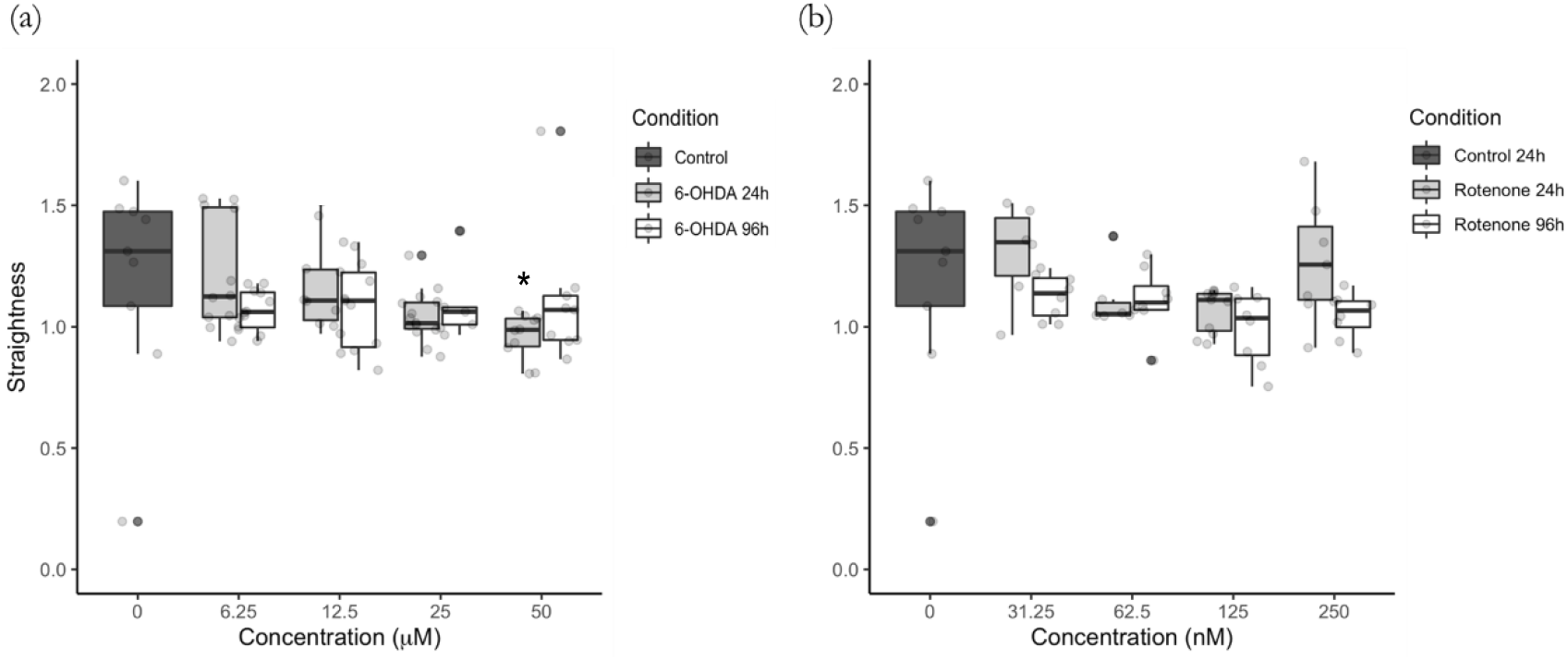
Mitochondrial movement pattern straightness was affected in cells treated with 6-OHDA (a) and rotenone (b). Mitochondria were labeled with the fluorescent dye MitoTracker Red CMXRos, their movement followed, and trajectory straightness was calculated as stated in Materials and Methods. Data are presented as boxplots in which each dot represents the mean of each mitochondria movement per video frame (n=5 to 15). Kruskal-Wallis test (One-way ANOVA on ranks) pair-wise (control vs 6-OHDA or control vs rotenone) was used to assess statistical significance, (*) p< 0.05.

Regarding individual mitochondria, the trajectory efficiency was small even in control cells (0.2 to 0.4) (Fig 6 a and b). Cells treated for 96 h with 6.25 μM, 12.5 μM, 25 μM and 50 μM 6-OHDA showed, on average, a decrease in 17%, 13%, 24% and 21%, respectively, in mitochondrial trajectory efficiency. Regarding cells treated for 24 h, the highest 6-OHDA concentration (50 μM) resulted in a 17% average decrease in mitochondria trajectory efficiency (Fig 6 a). Treatment with 62.5 nM for 24 h and with 125 nM rotenone (both 24 h and 96 h) displayed an average 15%, 21% and 21% decrease of mitochondrial trajectory efficiency when compared to control cells (Fig 6 b).

**Fig 6.**
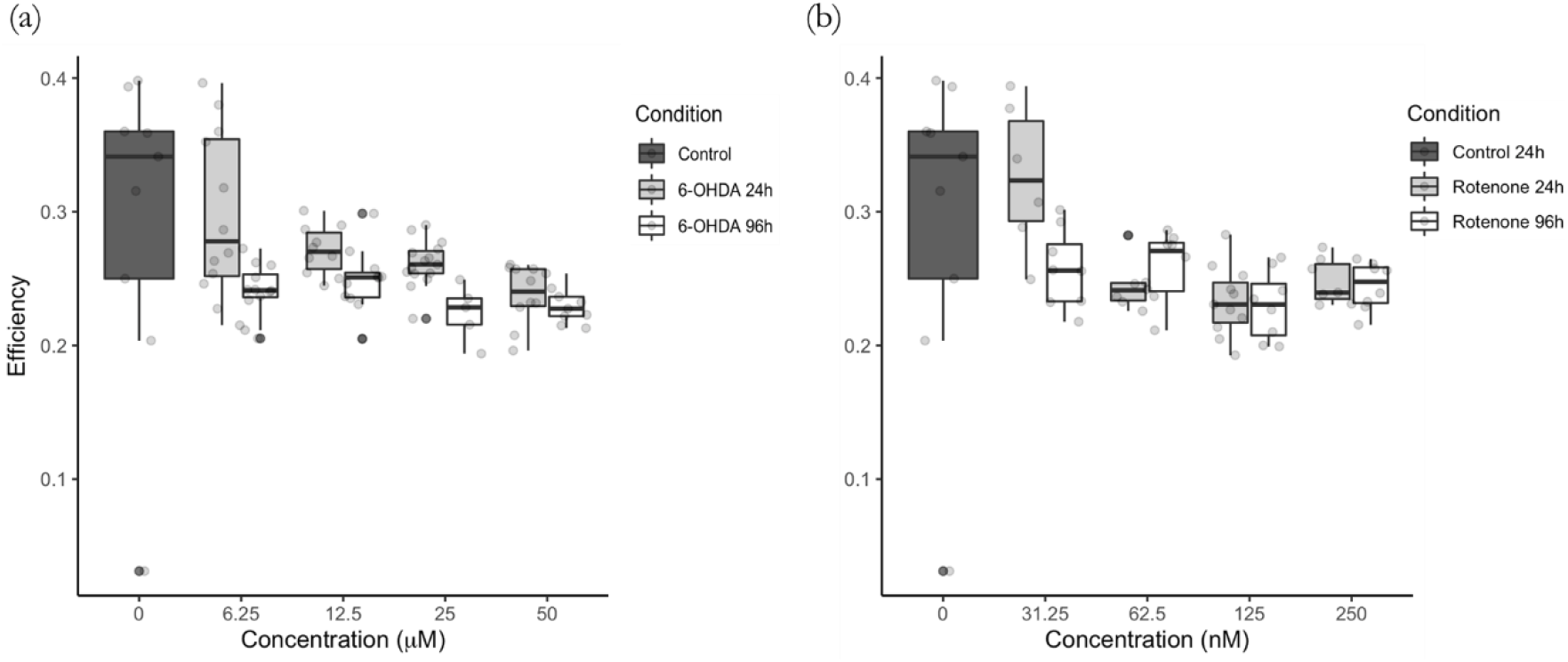
Mitochondrial trajectory efficiency was decreased in cells treated with 6-OHDA (a) and with rotenone (b). Mitochondria were labeled with the fluorescent dye MitoTracker Red CMXRos, their movement followed, and trajectory efficiency was calculated as stated in Materials and Methods. Data are represented as boxplots in which each dot represents the mean of each mitochondria movement per video frame (n=5 to 15). Kruskal-Wallis test (One-way ANOVA on ranks) pair-wise (control vs 6-OHDA or control vs rotenone) was used to assess statistical significance.

### 2.6. Mitochondria in cells treated with 6-OHDA and rotenone exhibit a higher degree of trajectory anisotropy

Cells incubated for 24 h and 96 h with 6-OHDA at all concentrations, with the exception of the highest concentration (50 μM) for both time points, showed a significant increase in mitochondrial trajectory anisotropy, which was reflected by more unidimensional trajectories (Fig 7 a). Regarding rotenone, the profile was different in 24 h treated cells. Mitochondria in cells incubated for 96 h with the lower rotenone concentrations, 31.25 nM and 62.5 nM, showed significantly higher degree of trajectory anisotropy, exhibiting a more unidimensional trajectory. However, cells treated for 24 h with rotenone at 62.5 nM and 250 nM of rotenone showed a significant elevation of the degree of mitochondrial trajectory anisotropy (Fig 7 b).

**Fig 7.**
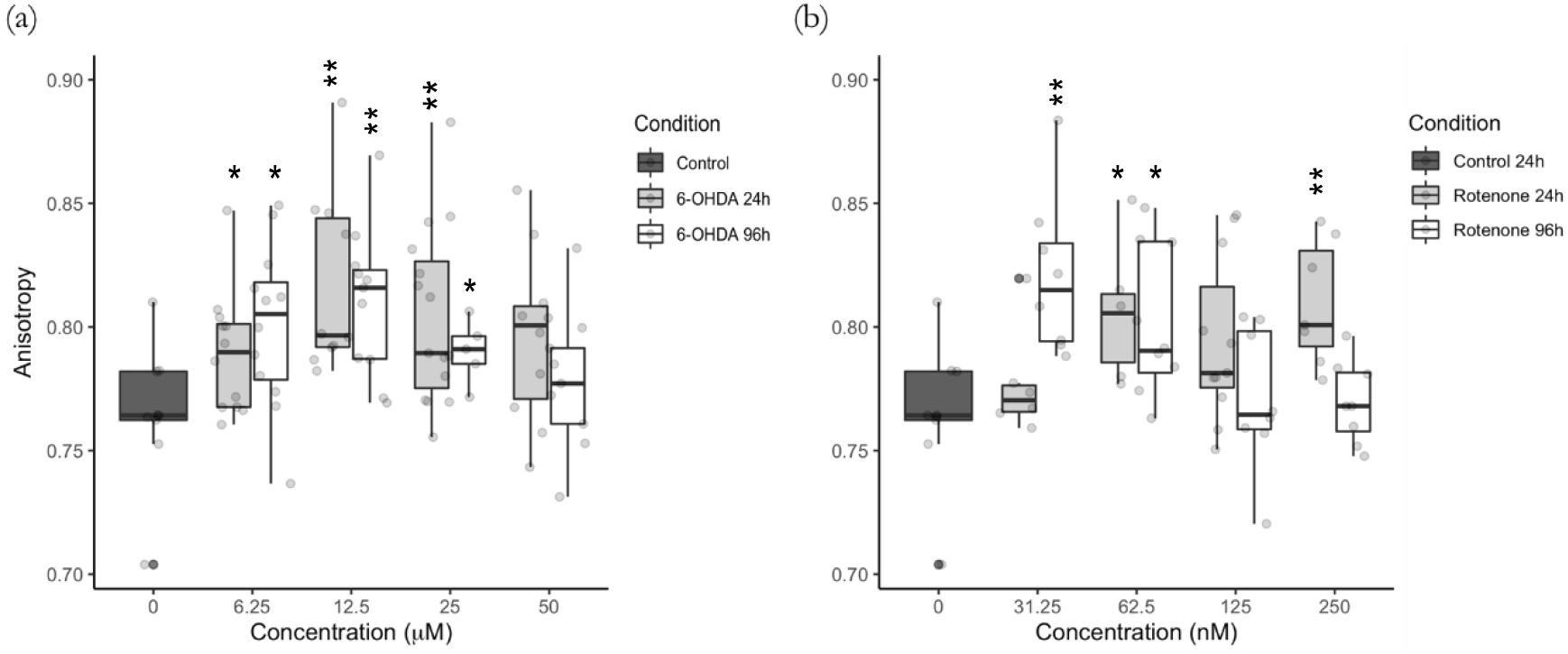
6-OHDA (a) and rotenone (b) promote higher degree of mitochondrial movement anisotropy. Mitochondria were labeled with the fluorescent dye MitoTracker Red CMXRos, their movement followed, and trajectory anisotropy was calculated as stated in Materials and Methods. Data are represented as boxplots in which each dot represents the mean of each mitochondria movement per video frame (n=5 to 15). Kruskal-Wallis test (One-way ANOVA on ranks) pair-wise (control vs 6-OHDA or control vs rotenone) was used to assess statistical significance, (**), p< 0.01, (*) p< 0.05.

### 2.7. Shorter incubation times with 6-OHDA enhance kurtosis of mitochondrial movement pattern

Significant increase of mitochondrial trajectory kurtosis was observed in cells treated for 24 h with the with 6-OHDA at 25 μM and 50 μM (Fig 8 a). On the other hand, no changes in kurtosis were observed for cells treated with 6-OHDA at the lower concentrations (Fig 8 a) and with rotenone at all concentrations (Fig 8 b).

**Fig 8.**
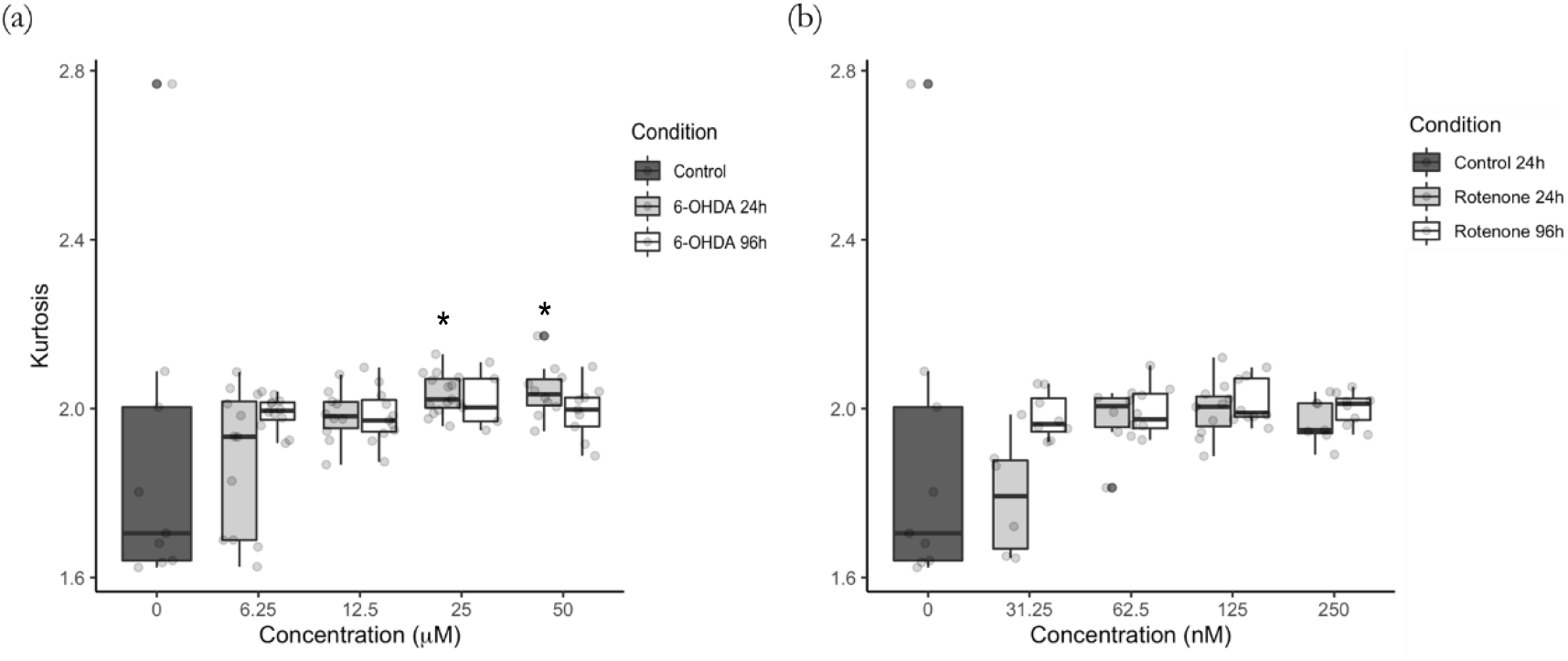
Kurtosis of mitochondrial movement and the effect of 6-OHDA (a) and rotenone (b). Mitochondria were labeled with the fluorescent dye MitoTracker Red CMXRos, their movement followed, and trajectory kurtosis was calculated as stated in Materials and Methods. Data are represented as boxplots in which each dot represents the mean of each mitochondria movement per video frame (n=5 to 15). Kruskal-Wallis test (One-way ANOVA on ranks) pair-wise (control vs 6-OHDA or control vs rotenone) was used to assess statistical significance, (*) p< 0.05.

### 2.8. Principal component analysis distinguishes control and treated cells

We performed PCA using the R stats library. The data were zero-centered and scaled to obtain unit variance before the analysis (z-score normalization) (31). In Fig 9, we show the PCA evaluation with the ellipses centered at the mean vector of the data points, which provide a visual intuition of the covariance (32). Treatments with 6-OHDA for 24 h presents a significant separation from the control (with variances of 62.2% and 37.8%), except for the lowest concentration (6.25 μM), at which the cluster shows a considerable superposition with control (Fig 9 a). The same pattern was observed for cells treated with rotenone for 24 h (with variances of 63.4% and 36.6%), in which the treatments data formed distinct clusters when compared to the control ones. However, the PCA shows essentially no difference between the control cells and those treated with 31.25 nM rotenone. For higher concentrations, a clockwise rotation of the principal axes with relation to control was observed, with higher variance along the anisotropy direction (Fig 9 b). In addition, for 96 h treatments with 6-OHDA (with variances 59.9% and 40.1%) (Fig 9 c) or rotenone (with variances 52.4% and 47.6%) (Fig 9 d), we observed that the results were grouped far from the control, indicating that a longer period of treatment may overcome the weak effect associated with lower concentrations. The data ellipses showed a trend for a covariance decrease as the concentration of 6-OHDA and rotenone increase for the 96 h treatments.

**Fig 9.**
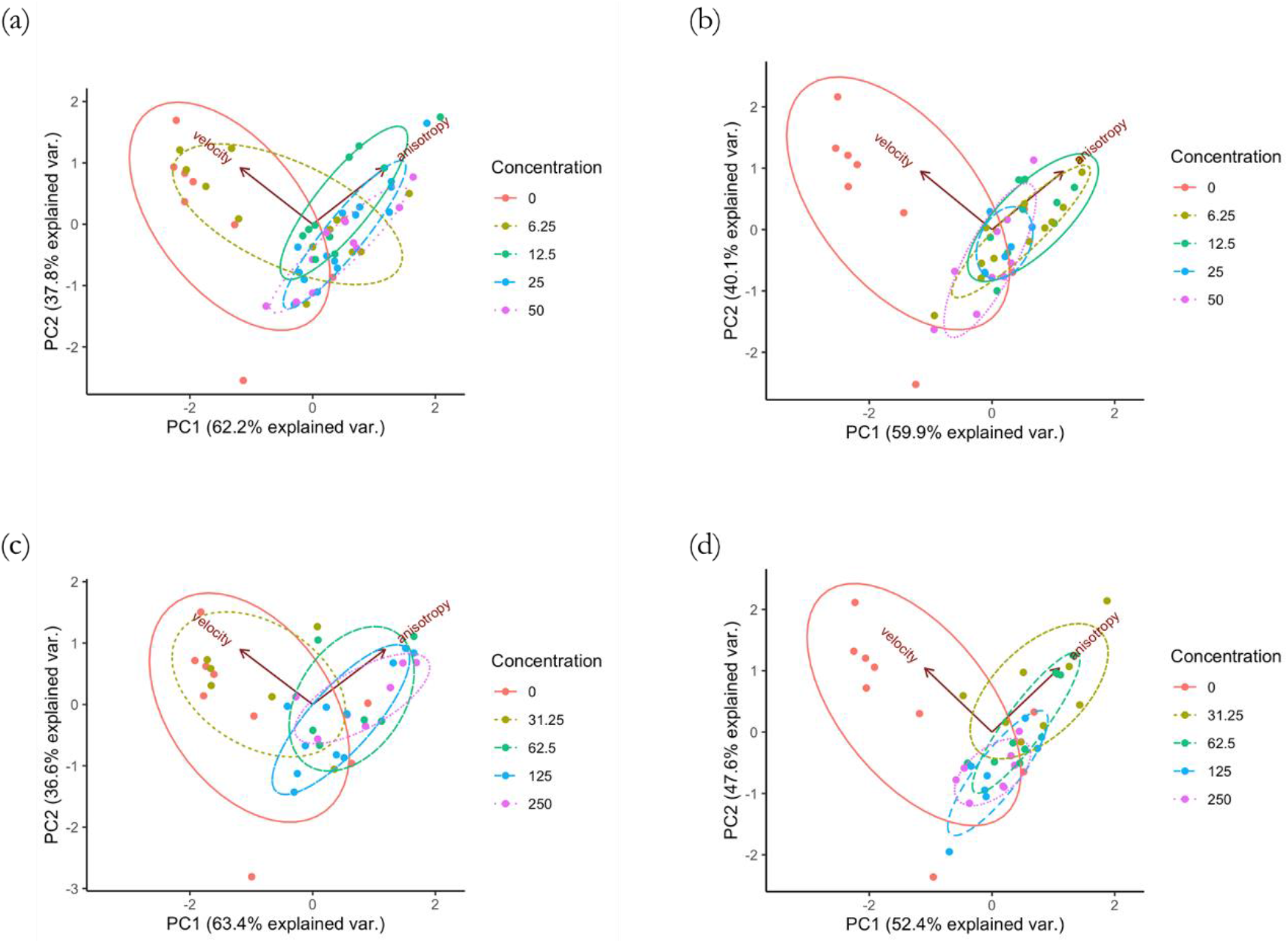
The panel shows PCAs for control together with each different treatment: 6-OHDA for 24 h (a) and 96 h (b) and rotenone for 24 h (c) and 96 h (d). The normal data ellipses are superposed. Only the two best features that better distinguish between the treatments and control conditions were considered, i.e. velocity and anisotropy. In general, the treatments grouped far from the control and presented higher variance along the anisotropy direction, apart from the treatments for 24 h with lowest concentration of 6-OHDA and rotenone.

## 3. Discussion

In recent years, the improvement of microscopy methods (enabling the acquisition of high signal-to-noise images) together with the development of automated particle tracking algorithms with certain level of accuracy allowed for the analysis of mitochondrial motility. However, a very careful and critical analysis should be performed when evaluating mitochondrial trafficking. A prime example is the study of mitochondrial mean velocity. It has been demonstrated that, depending on the cell model as well as method of mitochondrial tracking and movement analysis, the values of this parameter could range from an average of 0.1 μm/s to 1.5 μm/s (20, 21, 23, 33). To the best of our knowledge, no studies of mitochondrial trafficking have been performed using SH-SY5Y cells. Since we found that the rate of mitochondrial movement using these cells is 0.1−0.6 μm/s, SH-SY5Y cells present a plausible model for these studies.

In order to provide novel insights into mitochondria trajectory analysis and efficiency, we treated differentiated SH-SY5Y cells with 6-OHDA or rotenone and performed a more detailed analysis than carried out in previous studies presented in the literature. A key enhancement of our approach is the adoption of a wider range of features that help to characterize mitochondrial trajectories (such as their anisotropy, kurtosis, straightness and efficiency). A similar approach has been previously applied to the study of human natural killer cell migration in culture (34) and diffusion of nanoparticles in cellular microenvironment (35). 6-OHDA is a neurotoxic agent known to disrupt mitochondrial trafficking (36, 37). This substance is a hydroxylated analogue of the neurotransmitter dopamine (38) that induces mitochondrial toxicity by inhibiting complex I function, ensuing in superoxide production (39). 6-OHDA can also inhibit complex IV (40). Cells treated with 6-OHDA for 96 h exhibited a more indirect and less efficient trajectory featuring higher anisotropy. This may be explained by the negative synergistic effect of a significant decrease in ATP level and the level of βIII tubulin. Thus, 6-OHDA alters cell bioenergetics and microtubular tracks that are both indispensable for mitochondrial movement, ultimately resulting in a strong effect on dynamics of mitochondrial trafficking. Mitochondria trajectories in cells treated with this compound for 24 h presented, for higher concentrations (25 μM and 50 μM), a decrease in straightness, efficiency and net displacement but an increase in kurtosis. This weaker effect at shorter treatment times is possibly due to a smaller impact in reducing ATP levels.

Changes in mitochondrial trafficking have been described in cells treated with 6-OHDA. In a study using Lund human mesencephalic cells treated with 6-OHDA at 40 μM, 100 μM and 250 μM for 4 h and 7 h showed a decrease in the number of mitochondria moving both in the anterograde and retrograde direction without affecting the rate of mitochondrial movement (37). Similarly, it was shown that treatment with 60 μM 6-OHDA for 30 min in dopaminergic neurons decreased mitochondrial motility by approximately 50%. Again, no velocity alteration was evident under this scenario (36). Microtubule modifications and dynamics are also involved in 6-OHDA-related mitochondrial trafficking impairment. Related to our model, retinoic acid-differentiated SH-SY5Y cells treated with 30 μM 6-OHDA showed tubulin acetylation, which resulted in decreased microtubule growth rate, and increased level of monomeric tubulin, suggesting tubulin depolymerization. This effect was attributed to oxidative modifications of molecules of tubulin (41).

Rotenone is a time-dependent high-affinity irreversible inhibitor of complex I (42–44). This compound leads to inhibition of oxidative phosphorylation and oxygen consumption, ultimately triggering a cellular bioenergetic deficit. This agent induces oxidative damage of proteins, lipids and nucleic acids by means of generation of high levels of superoxide anion (45–47). Rotenone-treated cells showed weaker effect, when compared to their 6-OHDA-treated counterparts, when assessing the trajectory properties, which are a focus of this study. Although ATP levels and the level of βIII tubulin were decreased, no evident alterations were found in the trajectory straightness and kurtosis. Rotenone treatment, despite increasing anisotropy, indicated a more unidimensional trajectory and decreased trajectory efficiency.

Besides being involved in mitochondrial complex I inhibition, affecting ATP and superoxide anion production, neuronal cells treated with rotenone, both acutely and chronically, display alterations in mitochondrial trafficking. One study showed that primary cortical neurons acutely treated with 1 μM rotenone exhibited an increase in the number of stationary mitochondria. Additionally, a significant decrease in the mean velocity of mitochondrial movement in both directions was also reported (24). Using differentiated SH-SY5Y cells, it was shown that treatment of the cells with 50 nM rotenone for 8 and 16 days significantly suppressed the rate of mitochondrial trafficking. The authors hypothesized that the decrease in mitochondrial velocity was due to either the disruption of the microtubular network or oxidative stress (48). Indeed, several studies have shown that rotenone destabilizes microtubules, inducing tubulin depolymerization. Dopaminergic neurons incubated with 100 nM rotenone for 30 min displayed a significant increase in free tubulin (49). Additionally, incubation of cells with 10 μM rotenone for 12 h induced microtubule depolymerization and blocked its re-polymerization in a similar cell model (50). using non-neuronal cells, it was shown that rotenone induces tubulin conformational changes, affecting its secondary structure. This suppressed microtubule re-assembly and decreased the length of microtubules (51).

Examining one of the most frequently analyzed features of mitochondrial movement, which is the mean velocity of mitochondria along tubulin tracks, together with a rarely assessed feature of mitochondrial mobility, i.e. the anisotropy of mitochondrial trajectories, we were able to clearly distinguish between cells treated with neuronal poisons epitomized by 6-OHDA and rotenone. This was particularly evident at the longer treatment times of 96 h. By considering mean velocity and anisotropy combined with the PCA projection, we observed that the dispersion in velocity decreases with the treatment while for anisotropy increases. This behavior was observed in retinoic acid-differentiated SHSY5Y cells treated with both agents, also presenting a tendency for decreased variance in anisotropy for longer treatments (96 h) and for higher concentrations of 6-OHDA (50 μM) and rotenone (250 nM).

## 4. Conclusion

This study presents an innovative approach to quantitative analysis of mitochondria movement in differentiated SH-SY5Y cells treated with neuronal toxins at a range of concentrations and for different time points. Additionally to the conventionally studied movement characteristics such as mitochondrial net displacement and mean velocity, we introduced, for the first time, new movement descriptors to characterize mitochondria trajectories, i.e. their straightness, efficiency, anisotropy and kurtosis. We have demonstrated here for the first time that these new descriptors provide an insight into mitochondrial motility characteristics and can be used to characterize mitochondrial trajectories. Moreover, in cases in which mitochondrial length of movement and the movement duration, direction and velocity are not altered, these new trajectory descriptors can present a reliable and sensitive method to detect, in particular, the initial stages of neuronal degeneration.

## 5. Material and methods

### 5.1. Cell culture and treatments

SH-SY5Y cells (ECACC, cat. 94030304) were cultured in supplemented Dulbecco’s modified Eagle’s medium (DMEM, D5030, Sigma-Aldrich, USA) and differentiated into a neuronal-like morphology following a protocol published by us (52). Details are provided in the S1 Appendix.

### 5.2. ATP levels determination

Intracellular ATP was quantified using the CellTiter-Glo Luminescent Cell Viability Assay (G7570, Promega, USA) following manufacture’s protocol. Details are provided in the S1 Appendix.

### 5.3. Immunocytochemistry and fluorescence microscopy

βIII tubulin (sc80005, Santa Cruz, Germany) levels and Hoechst 33342 (B2261, Sigma-Aldrich) nuclear labelling in fixed SH-SY5Y cells were assessed following the protocol described in the S1 Appendix.

### 5.4. Live cell imaging

For live imaging, cells were differentiated in 35 mm μ-dishes (81156, Ibidi Germany) at 3×10^4^ cells/cm^2^ and treated with 6-OHDA and rotenone. Subsequently, mitochondria were stained with 25 nM of the mitochondrial fluorescent dye MitoTracker Red CMXRos (M7512, Invitrogen, Thermo Fisher Scientific) in the FluoroBrite DMEM Media (A1896702, Gibco, Thermo Fisher Scientific) for 30 min. The media was then replaced by fresh FluoroBrite DMEM Media. Movies of fluorescent mitochondria were then recorded using the TIRF-fitted Nikon Eclipse Ti2 inverted microscope. The lowest level of excitation light from the 561 nm laser was used for imaging, and the emitted light was collected using an mCherry filter. The EMCCD Andor iXon Ultra DU888 camera (Andor Technologies) was used to capture the images with resolution of 1024 × 1024 pixels (pixel size 13 × 13 μm) at 1 frame per second for 10 min.

#### Movie Processing

Raw movie files were convolved and filtered using ImageJ. After applying noise reduction, they were saved as a sequence of binary images. A MATLAB algorithm (www.github.com/kandelj/MitoSPT) (53) was then used to detect object movement across frames, allowing for the calculation of the trajectory, total and net distances traveled by each individual mitochondria. Movie processing details are provided in the S1 Appendix.

### 5.5. Quantitative analysis of trajectories

Specific physical properties, describing the curve shape and kinematics of individual mitochondria trajectories, were obtained with the python package trajpy (54, 55), available at https://github.com/ocbe-uio/trajpy/. Supplementary figures in S2 Appendix display some examples of trajectories. The calculated trajectory features are the following.

#### 5.5.1. Mean velocity

We evaluated the mitochondria mean velocity 〈*ν*〉 by calculating the ratio between the total length of the trajectory and the elapsed time Δ*t*

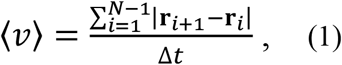

where *N* is the number of segments of the trajectory, and **r**_*i*_ is the position of the *i*-th point along the trajectory path.

#### 5.5.2. Anisotropy

The features related to the trajectory shape are functions of the gyration tensor obtained by the variance of the position along the trajectory. Mathematically, the components of the gyration tensor, *R_mn_*, are given by the following equation:

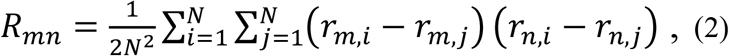

in which *m* and *n* are indices for the coordinates along the directions *x, y, z*.

Using the diagonalized gyration tensor *D* to define the tensor, 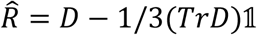 with the unity tensor 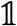, we obtained the degree of anisotropy among the principal axes (56), defined as

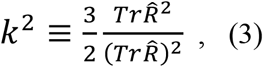

where, the setting 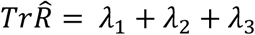, gives

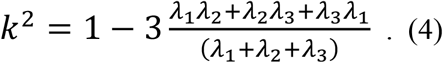

The minimum anisotropy, *k*^2^ = 0, is obtained when the distribution of the trajectory points is spherically symmetrical with λ_1_ = λ_2_ = λ_3_. The maximum anisotropy, *k*^2^ = 1, occurs when at least two eigenvalues are zero. High anisotropy refers to a small dimensionality in the principal axes coordinates - unidimensional trajectories present the highest anisotropy. Thus, anisotropy carries information about symmetry and dimensionality at the same time (57).

#### 5.5.3. Kurtosis

We obtained the kurtosis of the trajectory by projecting each position along the main principal eigenvector of the radius of the gyration tensor 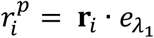, in which 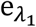 is the eigenvector associated to the eigenvalue λ_1_, and then calculating the quartic moment

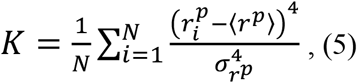

in which 〈*r^p^*〉 is the mean position of the projected trajectory and 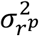 is its variance. Kurtosis is the measure of the ‘tailedness’ of the positions distribution in the trajectory (58).

#### 5.5.4. Straightness

Straightness compares the net displacement to the sum of displacements. It measures the likeliness of the trajectory to a straight line

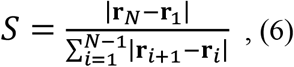

where **r**_1_ is the initial position and **r**_*N*_ is the last position on the trajectory. If the trajectory is completely straight, the numerator and denominator are the same, consequently *S* = 1. On the other hand, if 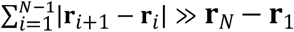 then *S* ≈ 0.

#### 5.5.5. Efficiency

Efficiency is similar to straightness described above. It is defined as the ratio between the net displacement and the sum of squared displacements:

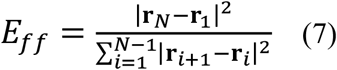

When a particle describes a long trajectory but ends at the same initial position, the measured efficiency will be zero. Moreover, for the same net displacement, a highly irregular trajectory will have smaller efficiency than the linear trajectory.

### 5.6. Statistics

Data were analyzed using R 4.0.3 (31). Results are presented in boxplots (box-and-whisker plots), in which the middle line represents the median and the whiskers go down to the minimum value and up to the maximum value, where each individual value is represented as a data point. The number of experiments carried out is presented in the legend of the figures.

We performed the non-parametric Kruskal-Wallis test pair-wise comparisons between the control and each treatment condition followed by Dunn’s post hoc for multiple conditions comparison. Statistical significance was set as (*) p<0.05, (**) p<0.01, (***) p<0.001 and (****) p<0.0001. Principal component analysis (PCA) was employed to identify the underlying covariable patterns of the data.

## Abbreviations

6-OHDA: 6-hydroxydopamine
ATP: adenosine triphosphate
BSA: bovine serum albumin
Ca^2+^: calcium
fps: frames per second
MIRO: mitochondrial rho
PCA: Principal Component Analysis
PBS: phosphate buffer saline
RA: retinoic Acid
TIRF: Total internal reflection fluorescence
TRAK: trafficking kinesin-binding.

## S1 Appendix

## Cell culture and differentiation

SH-SY5Y cells (ECACC, cat. 94030304) were cultured in Dulbecco’s modified Eagle’s medium (DMEM, D5030, Sigma-Aldrich, USA) containing 25 mM glucose (G7021, Sigma-Aldrich), 6 mM L-glutamine (G3126, Sigma-Aldrich), 5 mM HEPES (H4024, Sigma-Aldrich), 44 mM sodium bicarbonate (S6014, Sigma-Aldrich), 1 mM sodium pyruvate (P2256, Sigma-Aldrich), 10% (v/v) fetal bovine serum (41F6445K, Gibco, Thermo Fisher Scientific, USA) and 1% penicillin/streptomycin (1772652 Thermo Fisher Scientific) in a humidified atmosphere (5% CO_2_, 37 °C). Cell media was changed every 2 to 3 days, and cells were split when reaching 90-100% confluency.

For cell differentiation, cells were seeded at the density of 3×10^4^ cells/cm^2^ in low glucose (5 mM) media supplemented with 1% FBS and 10 μM retinoic acid (RA) (A6947 Panreac AppliChem ITW Reagents, Germany) for 3 days. Following differentiation, cells were treated with increasing concentration of 6-OHDA (H4381 Sigma-Aldrich) or rotenone (MKBS1062V, Sigma-Aldrich).

## ATP levels determination

Cell differentiation and treatments were accomplished in white, opaque-bottom, 96-well plates (136101, Thermo Fisher Scientific). At the end of cell treatments, the medium was removed and replaced by 50 μl of fresh medium. 50 μl of the Cell Titer-Glo reagent was added, and plates were agitated for 2 min on an orbital shaker to promote cell lysis. After 10-min incubation, the luminescent signal was recorded using Cytation™ 3 microplate reader (BioTek, USA).

## Immunocytochemistry and fluorescence microscopy

After cell differentiation and treatment, the cell culture medium was removed, cells were washed with warm phosphate buffer saline (PBS), fixed with 4% paraformaldehyde in PBS and stored at 4 °C. The cells were then washed 3 times with PBS and permeabilized with 0.2% (v/v) Triton X-100 (AC327371000, Fisher Scientific) in PBS for 2 min. The cells were then washed 3 times with PBS, and incubated with the blocking solution (3% bovine serum albumin, BSA; A6003 Sigma-Aldrich) in PBS. The cells were washed 3 times with PBS containing 1% BSA and incubated overnight at 4 °C with mouse anti-βIII tubulin (sc80005, Santa Cruz, Germany) at 1:200 dilution prepared in 3% BSA in PBS. This was followed by 90-min incubation with goat-anti-mouse Alexa Fluor 488 (A-11001, Cat. M7512, Invitrogen, Thermo Fisher Scientific, USA) at 1:1000 dilution in 3% BSA in PBS. Finally, cells were washed 3 times with 1% BSA in PBS and incubated with 1 μg/ml Hoechst 33342 (B2261, Sigma-Aldrich) in PBS for nuclei visualization.

Cell visualization was performed using an INCell Analyzer 2200 (GE Healthcare) cell imaging system. Images were acquired using a 20x objective (INCA ASAC 20 x/0.45, ELWD Plan Fluor). Image analysis was performed using the INCell Analyzer 1000 analysis software - Developer Toolbox. The image stack was uploaded by the software to identify our target set and to establish the respective parameters of area and number. The representative images shown in this work were visualized using ImageJ 1.52a (Wayne Rasband, National Instituted of Health, USA).

## ImageJ image pre-processing

Following the published protocol (53), we pre-processed raw image files using ImageJ. Briefly, time-lapse images were first convolved using the 5×5 edge-detection, converted to the frequency domain using a Fast Fourier Transform, and then subjected to a bandpass filter ranging from 2 pixels (~0.3 μm) to 100 pixels (~16 μm). The resulting images were manually thresholded to eliminate the noise, and the results saved as a sequence of individual binary images.

## MATLAB Algorithm

The stacks of individual images were analyzed by an open source MATLAB algorithm (www.github.com/kandelj/MitoSPT) (53). Briefly, the algorithm read each frame into MATLAB and used the built-in functions *bwconncomp* and *regionprops* to find the connected white objects and to measure their sizes, respectively. The image was then recreated to contain only objects with the area within the specified limits defined by the user. Each frame went through the same process. The current frame objects were labeled or re-labeled by comparing their pixel locations with the ones from the previous frame. After all objects were labeled/re-labeled, their locations were stored, and they were prepared to be compared with the next frame. After this process was repeated frame by frame, the collected centroid locations were used to calculate the total and net distances traveled by each object (53). In addition, the software was adapted to output the raw trajectories of each individual mitochondria into a comma-separated values file (csv) for external analysis.

## S2 Appendix

## Supplementary figures captions

**Supplementary Fig. 1** – A subset of 3 trajectories obtained from the control group.

**Supplementary Fig. 2** – Three trajectories with different stochastic noise strength *γ* = {0,6,20}. This example is a linear trajectory *y(x)* = *x* + *γ*(Rand − 0.5) under the influence of a random noise with {*x* ∊ ℝ | 0 ≤ *x* ≤ 100} and *γ* is the parameter that controls the stochastic strength. It is shown the trajectories for *γ* = 0 (without noise), *γ* = 6 (weak noise) and (high noise), presenting lower to higher tortuosity, respectively.

**Supplementary Fig. 3** – Circular trajectory to observe the effect of symmetry in the features by considering subsets of the circle, as exemplified with 6, 11 and 20 points. We calculated the anisotropy, kurtosis, straightness and efficiency attributes for incomplete circles from 3 to 20 points (complete circle), counterclockwise, and determined the dependency as a function of the number of points considered.

**Supplementary Fig. 4** – The effect of stochasticity on each of the features is depicted. We can observe that anisotropy and kurtosis are resilient to the introduction of stochastic noise in the trajectory. The anisotropy shows a tendency to decrease as the noise influence increases, while the kurtosis goes in the opposite direction and increases with *γ*. In contrast, efficiency and straightness are strongly affected by stochasticity, decreasing rapidly.

**Supplementary Fig. 5** – Anisotropy, kurtosis, efficiency and straightness measured for the circular trajectory with different subsets. We can see that the anisotropy and the kurtosis present a non-monotonic behavior. As we consider more points in the circle, the anisotropy decreases due to the symmetry of the circle. With 11 points we have the semi-circle, which coincides with a local minimum in anisotropy and a local maximum in kurtosis. Efficiency and straightness decrease monotonically as we vary the number of points. The examples explored here highlight the difficulties faced in the analysis of some features, presenting often a non-intuitive behavior.

## References

1. Sheng ZH, Cai Q. Mitochondrial transport in neurons: impact on synaptic homeostasis and neurodegeneration. Nat Rev Neurosci. 2012;13(2):77–93.

2. Saxton WM, Hollenbeck PJ. The axonal transport of mitochondria. J Cell Sci. 2012;125(Pt 9):2095–104.

3. Sheng ZH. Mitochondrial trafficking and anchoring in neurons: New insight and implications. J Cell Biol. 2014;204(7):1087–98.

4. Attwell D, Laughlin SB. An energy budget for signaling in the grey matter of the brain. J Cereb Blood Flow Metab. 2001;21(10):1133–45.

5. Harris JJ, Jolivet R, Attwell D. Synaptic energy use and supply. Neuron. 2012;75(5):762–77.

6. Sun T, Qiao H, Pan PY, Chen Y, Sheng ZH. Motile axonal mitochondria contribute to the variability of presynaptic strength. Cell Rep. 2013;4(3):413–9.

7. Billups B, Forsythe ID. Presynaptic mitochondrial calcium sequestration influences transmission at mammalian central synapses. J Neurosci. 2002;22(14):5840–7.

8. Medler K, Gleason EL. Mitochondrial Ca(2+) buffering regulates synaptic transmission between retinal amacrine cells. J Neurophysiol. 2002;87(3):1426–39.

9. Schwarz TL. Mitochondrial trafficking in neurons. Cold Spring Harb Perspect Biol. 2013;5(6).

10. Morris RL, Hollenbeck PJ. The regulation of bidirectional mitochondrial transport is coordinated with axonal outgrowth. J Cell Sci. 1993;104 (Pt 3):917–27.

11. Pilling AD, Horiuchi D, Lively CM, Saxton WM. Kinesin-1 and Dynein are the primary motors for fast transport of mitochondria in Drosophila motor axons. Mol Biol Cell. 2006;17(4):2057–68.

12. Hirokawa N, Niwa S, Tanaka Y. Molecular motors in neurons: transport mechanisms and roles in brain function, development, and disease. Neuron. 2010;68(4):610–38.

13. Kapitein LC, Hoogenraad CC. Building the Neuronal Microtubule Cytoskeleton. Neuron. 2015;87(3):492–506.

14. Nguyen MM, Stone MC, Rolls MM. Microtubules are organized independently of the centrosome in Drosophila neurons. Neural Dev. 2011;6:38.

15. Yau KW, Schatzle P, Tortosa E, Pages S, Holtmaat A, Kapitein LC, et al. Dendrites In Vitro and In Vivo Contain Microtubules of Opposite Polarity and Axon Formation Correlates with Uniform Plus-End-Out Microtubule Orientation. J Neurosci. 2016;36(4):1071–85.

16. MacAskill AF, Brickley K, Stephenson FA, Kittler JT. GTPase dependent recruitment of Grif-1 by Miro1 regulates mitochondrial trafficking in hippocampal neurons. Mol Cell Neurosci. 2009;40(3):301–12.

17. Brickley K, Stephenson FA. Trafficking kinesin protein (TRAK)-mediated transport of mitochondria in axons of hippocampal neurons. J Biol Chem. 2011;286(20):18079–92.

18. King SJ, Schroer TA. Dynactin increases the processivity of the cytoplasmic dynein motor. Nat Cell Biol. 2000;2(1):20–4.

19. Ligon LA, Steward O. Movement of mitochondria in the axons and dendrites of cultured hippocampal neurons. J Comp Neurol. 2000;427(3):340–50.

20. Misgeld T, Kerschensteiner M, Bareyre FM, Burgess RW, Lichtman JW. Imaging axonal transport of mitochondria in vivo. Nat Methods. 2007;4(7):559–61.

21. Fang C, Bourdette D, Banker G. Oxidative stress inhibits axonal transport: implications for neurodegenerative diseases. Mol Neurodegener. 2012;7:29.

22. Bros H, Millward JM, Paul F, Niesner R, Infante-Duarte C. Oxidative damage to mitochondria at the nodes of Ranvier precedes axon degeneration in ex vivo transected axons. Exp Neurol. 2014;261:127–35.

23. Bros H, Hauser A, Paul F, Niesner R, Infante-Duarte C. Assessing Mitochondrial Movement Within Neurons: Manual Versus Automated Tracking Methods. Traffic. 2015;16(8):906–17.

24. Chen M, Li Y, Yang M, Chen X, Chen Y, Yang F, et al. A new method for quantifying mitochondrial axonal transport. Protein Cell. 2016;7(11):804–19.

25. Coutu DL, Schroeder T. Probing cellular processes by long-term live imaging--historic problems and current solutions. J Cell Sci. 2013;126(Pt 17):3805–15.

26. Gerencser AA, Nicholls DG. Measurement of instantaneous velocity vectors of organelle transport: mitochondrial transport and bioenergetics in hippocampal neurons. Biophys J. 2008;95(6):3079–99.

27. Chang DT, Rintoul GL, Pandipati S, Reynolds IJ. Mutant huntingtin aggregates impair mitochondrial movement and trafficking in cortical neurons. Neurobiol Dis. 2006;22(2):388–400.

28. Axelrod D. Chapter 7: Total internal reflection fluorescence microscopy. Methods Cell Biol. 2008;89:169–221.

29. Mattheyses AL, Simon SM, Rappoport JZ. Imaging with total internal reflection fluorescence microscopy for the cell biologist. J Cell Sci. 2010;123(Pt 21):3621–8.

30. Poulter NS, Pitkeathly WT, Smith PJ, Rappoport JZ. The physical basis of total internal reflection fluorescence (TIRF) microscopy and its cellular applications. Methods Mol Biol. 2015;1251:1–23.

31. R Core Team. R: A language and environment for statistical computing. R Foundation for Statistical Computing. Vienna, Austria. 2020 [Available from: http://www.r-project.org/index.html.

32. Friendly M, Monette G, Fox J. Elliptical Insights: Understanding Statistical Methods through Elliptical Geometry. Statist Sci. 2013;28(1):1–39.

33. MacAskill AF, Kittler JT. Control of mitochondrial transport and localization in neurons. Trends Cell Biol. 2010;20(2):102–12.

34. Lee BJ, Mace EM. Acquisition of cell migration defines NK cell differentiation from hematopoietic stem cell precursors. Mol Biol Cell. 2017;28(25):3573–81.

35. Wagner T, Kroll A, Haramagatti CR, Lipinski HG, Wiemann M. Classification and Segmentation of Nanoparticle Diffusion Trajectories in Cellular Micro Environments. PLoS One. 2017;12(1):e0170165.

36. Lu X, Kim-Han JS, Harmon S, Sakiyama-Elbert SE, O’Malley KL. The Parkinsonian mimetic, 6-OHDA, impairs axonal transport in dopaminergic axons. Mol Neurodegener. 2014;9:17.

37. Stepkowski TM, Meczynska-Wielgosz S, Kruszewski M. mitoLUHMES: An Engineered Neuronal Cell Line for the Analysis of the Motility of Mitochondria. Cell Mol Neurobiol. 2017;37(6):1055–66.

38. Blum D, Torch S, Lambeng N, Nissou M, Benabid AL, Sadoul R, et al. Molecular pathways involved in the neurotoxicity of 6-OHDA, dopamine and MPTP: contribution to the apoptotic theory in Parkinson’s disease. Prog Neurobiol. 2001;65(2):135–72.

39. Betarbet R, Sherer TB, Greenamyre JT. Animal models of Parkinson’s disease. Bioessays. 2002;24(4):308–18.

40. Glinka YY, Youdim MBH. Inhibition of mitochondrial complexes I and IV by 6-hydroxydopamine. European Journal of Pharmacology: Environmental Toxicology and Pharmacology. 1995;292(3):329–32.

41. Patel VP, Defranco DB, Chu CT. Altered transcription factor trafficking in oxidatively-stressed neuronal cells. Biochim Biophys Acta. 2012;1822(11):1773–82.

42. Schuler F, Casida JE. Functional coupling of PSST and ND1 subunits in NADH:ubiquinone oxidoreductase established by photoaffinity labeling. Biochim Biophys Acta. 2001;1506(1):79–87.

43. Degli Esposti M. Inhibitors of NADH-ubiquinone reductase: an overview. Biochim Biophys Acta. 1998;1364(2):222–35.

44. Lummen P. Complex I inhibitors as insecticides and acaricides. Biochim Biophys Acta. 1998;1364(2):287–96.

45. Sanders LH, Timothy Greenamyre J. Oxidative damage to macromolecules in human Parkinson disease and the rotenone model. Free Radic Biol Med. 2013;62:111–20.

46. Sherer TB, Betarbet R, Testa CM, Seo BB, Richardson JR, Kim JH, et al. Mechanism of toxicity in rotenone models of Parkinson’s disease. J Neurosci. 2003;23(34):10756–64.

47. Uversky VN. Neurotoxicant-induced animal models of Parkinson’s disease: understanding the role of rotenone, maneb and paraquat in neurodegeneration. Cell Tissue Res. 2004;318(1):225–41.

48. Borland MK, Trimmer PA, Rubinstein JD, Keeney PM, Mohanakumar K, Liu L, et al. Chronic, low-dose rotenone reproduces Lewy neurites found in early stages of Parkinson’s disease, reduces mitochondrial movement and slowly kills differentiated SH-SY5Y neural cells. Mol Neurodegener. 2008;3:21.

49. Jiang Q, Yan Z, Feng J. Neurotrophic factors stabilize microtubules and protect against rotenone toxicity on dopaminergic neurons. J Biol Chem. 2006;281(39):29391–400.

50. Ren Y, Liu W, Jiang H, Jiang Q, Feng J. Selective vulnerability of dopaminergic neurons to microtubule depolymerization. J Biol Chem. 2005;280(40):34105–12.

51. Srivastava AS, Feng Z, Mishra R, Malhotra R, Kim HS, Carrier E. Embryonic stem cells ameliorate piroxicam-induced colitis in IL10-/-KO mice. Biochem Biophys Res Commun. 2007;361(4):953–9.

52. Simoes RF, Ferrao R, Silva MR, Pinho SLC, Ferreira L, Oliveira PJ, et al. Refinement of a differentiation protocol using neuroblastoma SH-SY5Y cells for use in neurotoxicology research. Food Chem Toxicol. 2021;149:111967.

53. Kandel J, Chou P, Eckmann DM. Automated detection of whole-cell mitochondrial motility and its dependence on cytoarchitectural integrity. Biotechnol Bioeng. 2015;112(7):1395–405.

54. Moreira-Soares M. trajpy. 1.3.1 ed: Zenode; 2020.

55. Moreira-Soares M, Cunha SP, Bordin JR, Travasso RDM. Adhesion modulates cell morphology and migration within dense fibrous networks. Journal of Physics: Condensed Matter. 2020;32(31):314001.

56. Theodorou DN, Suter UW. Shape of unperturbed linear polymers: polypropylene. Macromolecules. 1985;18(6):1206–14.

57. Arkin H, Janke W. Gyration tensor based analysis of the shapes of polymer chains in an attractive spherical cage. J Chem Phys. 2013;138(5):054904.

58. Helmuth JA, Burckhardt CJ, Koumoutsakos P, Greber UF, Sbalzarini IF. A novel supervised trajectory segmentation algorithm identifies distinct types of human adenovirus motion in host cells. J Struct Biol. 2007;159(3):347–58.

