## Supplementary material for "Quantitative analysis of neuronal mitochondrial movement reveals patterns resulting from neurotoxicity of rotenone and 6-hydroxydopamine": S2 Appendix

**Supplementary figures**

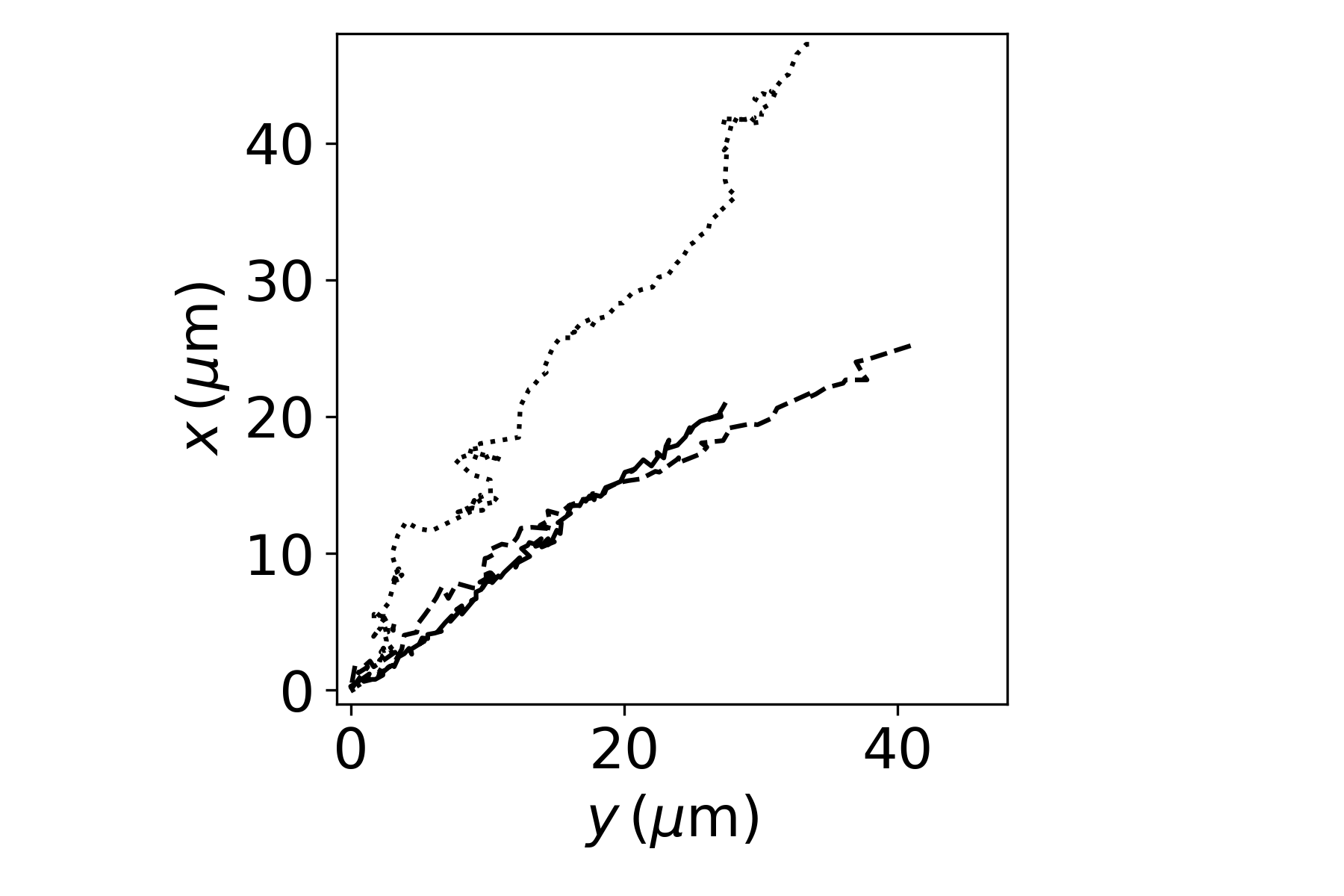

**Supplementary Fig. 1 -** A subset of 3 trajectories obtained from the control group.

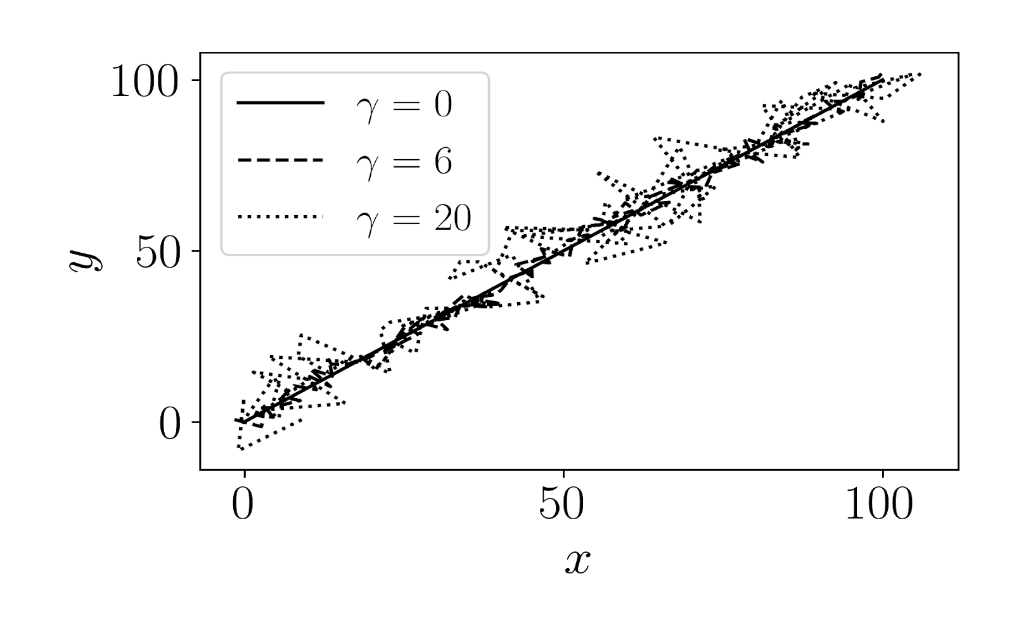

**Supplementary Fig. 2** - Three trajectories with different stochastic noise strength
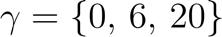
<https://www.codecogs.com/eqnedit.php?latex=%5Cgamma%20=%20%5C%7B0,%20%5C,%206,%5C,%2020%5C%7D%230> . This example is a linear trajectory
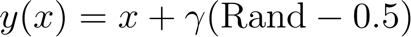
<https://www.codecogs.com/eqnedit.php?latex=y(x)%20=%20x%20+%20%5Cgamma%20(%5Cmathrm%7BRand%7D-0.5)%20%230> under the influence of a random noise with
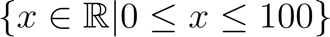
<https://www.codecogs.com/eqnedit.php?latex=%20%5C%7B%20x%20%5Cin%20%5Cmathbb%7BR%7D%20%7C%200%20%5Cle%20x%20%5Cle%20100%20%5C%7D%230> and
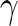
<https://www.codecogs.com/eqnedit.php?latex=%5Cgamma%230> is the parameter that controls the stochastic strength. It is shown the trajectories for
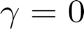
<https://www.codecogs.com/eqnedit.php?latex=%5Cgamma%20=%200%20%230> (without noise),
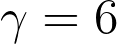
 (weak noise) and
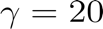
<https://www.codecogs.com/eqnedit.php?latex=%5Cgamma%20=%2020%230> (high noise), presenting lower to higher tortuosity, respectively.

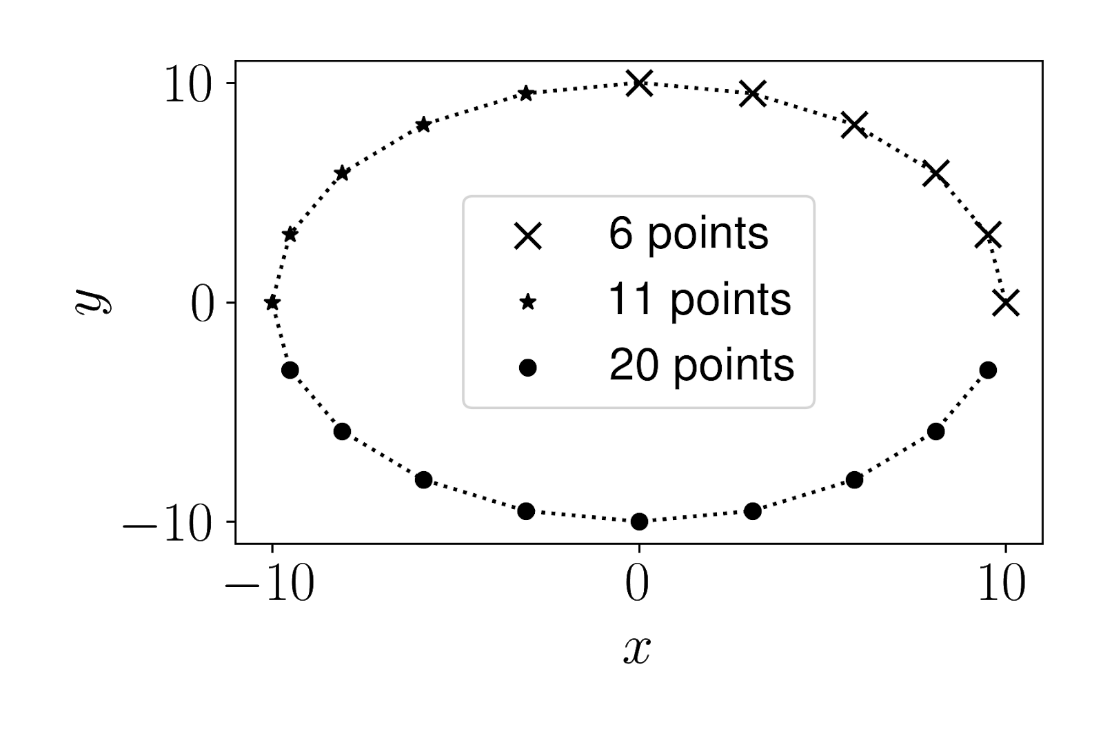

**Supplementary Fig. 3** - Circular trajectory to observe the effect of symmetry in the features by considering subsets of the circle, as exemplified with 6, 11 and 20 points. We calculated the anisotropy, kurtosis, straightness and efficiency attributes for incomplete circles from 3 to 20 points (complete circle), counterclockwise, and determined the dependency as a function of the number of points considered.

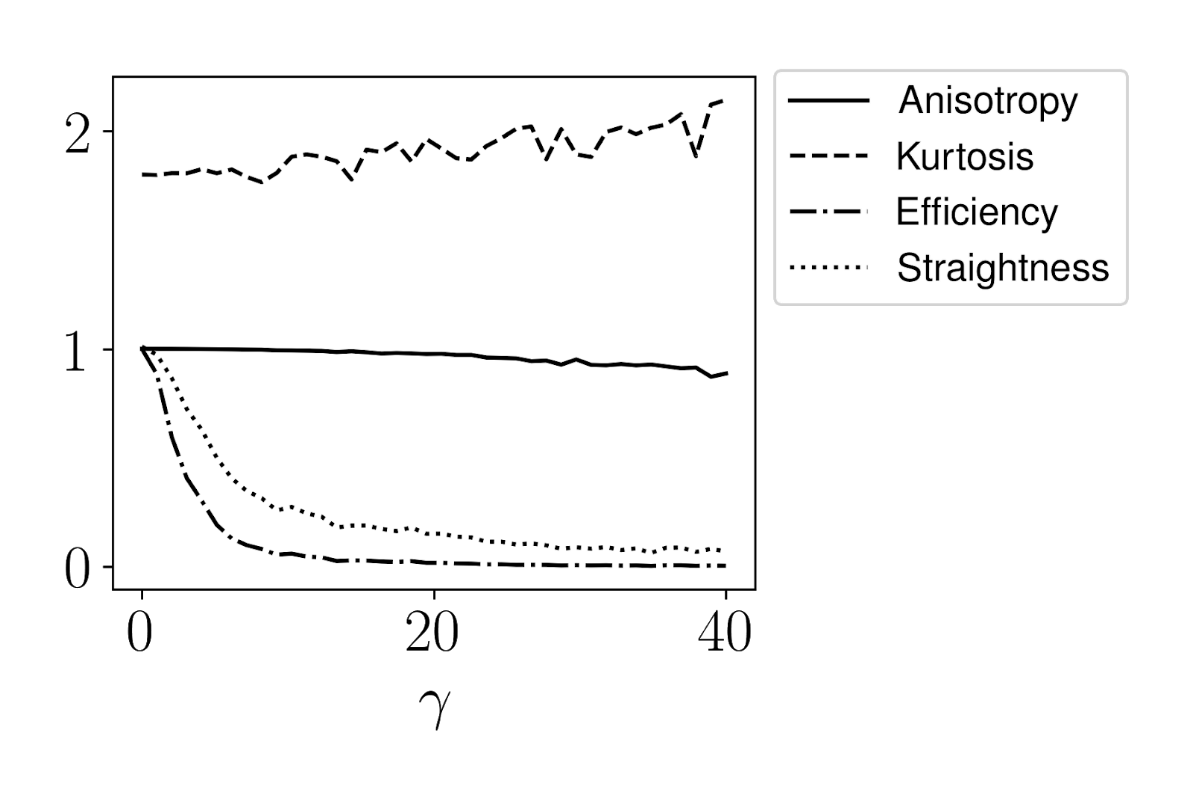

**Supplementary Fig. 4** - The effect of stochasticity on each of the features is depicted. We can observe that anisotropy and kurtosis are resilient to the introduction of stochastic noise in the trajectory. The anisotropy shows a tendency to decrease as the noise influence increases, while the kurtosis goes in the opposite direction and increases with
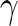
<https://www.codecogs.com/eqnedit.php?latex=%5Cgamma%230>. In contrast, efficiency and straightness are strongly affected by stochasticity, decreasing rapidly.

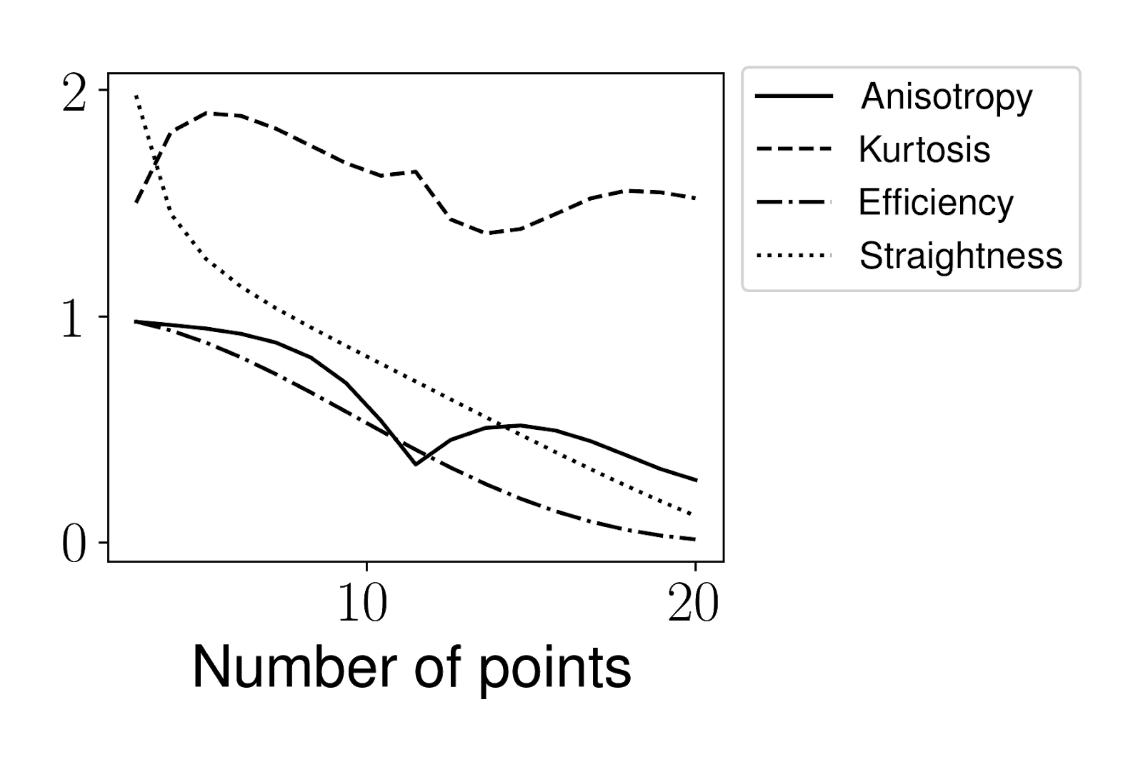

**Supplementary Fig. 5** - Anisotropy, kurtosis, efficiency and straightness measured for the circular trajectory with different subsets. We can see that the anisotropy and the kurtosis present a non-monotonic behavior. As we consider more points in the circle, the anisotropy decreases due to the symmetry of the circle. With 11 points we have the semi-circle, which coincides with a local minimum in anisotropy and a local maximum in kurtosis. Efficiency and straightness decrease monotonically as we vary the number of points. The examples explored here highlight the difficulties faced in the analysis of some features, presenting often a non-intuitive behavior.
